# Novel ddPCR diagnostic assays for the high-throughput surveillance of *Kdr* mutations in *Aedes albopictus*

**DOI:** 10.64898/2026.08.06.743222

**Authors:** Louis Nadalin, Thierry Gaude, Frédéric Laporte, Julien Renaud, Claudia Mulat, Delphine Rey, Grégory L’ambert, Nicolas Le Doeuff-Le Roy, Guillaume Lacour, Antoine Mignotte, Thomas Althaus, Alizée Costantini, Konstantinos Mavridis, Verena Pichler, Beniamino Caputo, Jean-Marc Bonneville, Jean-Philippe David

## Abstract

**Background:** Resistance of the arbovirus vector *Aedes albopictus* to pyrethroid insecticides is an emerging threat to vector control programs in Europe. Three knock-down resistance (’*Kdr*’) mutations affecting the voltage-gated sodium channels targeted by pyrethroids are known to confer resistance: *V1016G, I1532T* and *F1534C*. As these *Kdr* mutations are actively circulating in Europe, their monitoring is crucial for resistance surveillance programs. However, current *Kdr* genotyping methods are labor-intensive and costly, limiting their applicability for large-scale high-throughput surveillance.

**Methodology:** Novel digital droplet PCR (ddPCR) TaqMan assays were developed, allowing the quantification of these *Kdr* mutations from pooled mosquito samples from field populations. The specificity of these assays was validated by using mosquitoes of known genotypes together with synthetic DNA constructs carrying haplotype combinations previously unseen in the field. The assays’ accuracy was assessed by comparing *Kdr* frequencies measured from pooled mosquitoes to those derived from individual genotypes. These assays were then used in a pilot surveillance study in mainland France, integrating deltamethrin bioassays, pooled *Kdr* mutation tracking and vector control interventions data.

**Findings:** The developed ddPCR assays demonstrated high accuracy and specificity, with matching *Kdr* mutation frequencies between pooled samples and individual genotyping. The pilot surveillance study confirmed the low prevalence of *Kdr V1016G* and *I1532T* mutations in most French populations, though some populations exhibited a moderate decrease in susceptibility to deltamethrin. Deltamethrin spraying intensity was weakly correlated with *Kdr I1532T* mutation frequency while no correlation was observed with deltamethrin susceptibility, suggesting that insecticide selective pressure from curative vector control activities is at most a minor driver of resistance.

**Conclusion:** These novel ddPCR assays provide a simple, cost-effective and high-throughput method for quantifying the frequency of *Kdr* mutations in *Ae. albopictus* populations. Their implementation in routine country-wide surveillance programs will enhance the detection of emerging resistance, and inform vector control strategies, preventing arboviral disease transmission.

**Author summary:** *Aedes albopictus*, the Asian tiger mosquito, is an emerging global threat due to its ability to transmit the dengue, chikungunya and Zika viruses among others. Recurrent use of insecticides to prevent infections selects resistant mosquitoes. *Kdr* (knock-down resistance) mutations confer resistance to pyrethroid insecticides, like deltamethrin. In order to track the arrival of these mutations in a population, individual PCR tests are routinely used, which is very inefficient and costly. The digital droplet PCR tests presented in this study can instead be performed on pools of mosquitoes from the field, greatly increasing the output while reducing the costs. This makes them ideal for use in the surveillance of these mutations in field populations. We first confirmed that the tests are as efficient in pools as they are on single mosquitoes, then we applied these tests in a pilot study on French field mosquitoes to demonstrate their applicability for use in surveillance. Finally, we explored the link between deltamethrin sprayings, bioassay mortality and *Kdr* mutations frequency, which is non-existent. This means the current management of mosquito populations via insecticide use is sensible. We believe these tests have a place in streamlining the future surveillance programs of these mutations in the field.

## Introduction

Vector-borne diseases account for more than 700 000 deaths per annum worldwide, making them a major public health concern [1]. *Aedes albopictus*, the Asian tiger mosquito, is vector of several human arboviruses such as dengue, chikungunya and Zika. As of April 2026, 16 member states of the European Union (EU) have recorded long-term *Aedes albopictus* establishment, including mainland France [2]. Such establishment in the EU has led to an increased rate of arbovirus transmission [3, 4]: for instance, mainland France has seen 809 indigenous chikungunya cases in 2025, its highest number recorded [5]. In order to prevent outbreaks, vector control interventions aiming at reducing mosquito populations are routinely carried out in EU countries and frequently include spatial sprayings with pyrethroid insecticides in high risk areas, or in the vicinity of detected arbovirus cases [6].

Insecticide resistance is an adaptation of insect populations to a repeated insecticide pressure [7]. For mosquitoes, this selective pressure result from vector control activities and the use of insecticides at home or in agriculture [8–10]. Pyrethroid insecticides, such as deltamethrin, are among the most used insecticides globally [11]. They bind to the voltage-gated sodium channels (VGSC), leading to paralysis and subsequent death of the insect [12]. However, as seen in other mosquito species, the recurrent use of pyrethroids against *Ae. albopictus* populations leads to the selection of pyrethroid resistance mechanisms [13]. Among these, Knock-down resistance (‘*Kdr* ‘) mutations affecting the VGSCs are known to confer high resistance to most pyrethroids [14]. In contrast to *Ae. aegypti*, in which decades of pyrethroid usage led to the emergence of multiple *Kdr* mutations, often combined as haplotypes [15], only a few *Kdr* mutations have yet been associated with pyrethroid resistance in *Ae. albopictus* [16]. These include the *V1016G, I1532T* and *F1534C* mutations that have likely been introduced in southern Europe from Asia.

The *V1016G* mutation was first identified in Italy [17] and then observed in other EU countries such as Spain and mainland France [18, 19]. The *I1532T* mutation was also detected in Italy, mainland France and Greece [19–21]. The *F1534C* mutation was originally identified in Singapore [22] and then detected in western EU, including Spain [23, 24]. This makes southern Europe a potential hotspot for the emergence of pyrethroid resistance in *Ae. albopictus* and its spread further inland in the EU.

Considering the rapid spread of *Ae. albopictus* in Europe and the intensification of pyrethroid usage for arbovirus and nuisance control, the surveillance of resistance has become a major issue in sustaining the efficacy of vector control, deltamethrin being the only adulticide used to prevent arboviral transmission in France. The surveillance of phenotypic resistance can be achieved by performing regular insecticide bioassays, though this is extremely laborious, requires dedicated staff and infrastructure, and does not inform about resistance mechanisms [25]. By contrast, molecular assays allow tracking known resistance alleles at a high throughput for limited cost and effort. Some PCR-based assays have been developed to track *Kdr* mutations *Ae. albopictus* [26, 27]. These genotyping assays use single mosquito specimens and thus require performing dozens of gDNA extractions and PCR reactions per mosquito population, leading to significant costs and labwork at a country-wide scale. Multiplexed mass sequencing of PCR products can also be used to efficiently track *Kdr* mutations at a country scale but such approaches still require individual sample processing and complex downstream bioinformatic analyses [19]. Considering that *Kdr* mutation frequencies are sufficient to track resistance, performing surveillance at the population level (*i*.*e*. using a single gDNA extraction and PCR reaction per population) would represent a significant throughput and cost improvement for the routine surveillance of resistance at the country level.

Digital droplet PCR (ddPCR) is an emulsion-based PCR technology that allows performing thousands of PCR reactions within a single PCR well [28]. This fractioning of the PCR reaction allows for a precise quantification of the DNA target [29]. As a consequence, this technology provides the opportunity to directly quantify the frequency of *Kdr* alleles from pools of mosquitoes using a single DNA extraction and PCR amplification per population.

In this context, the present study aimed at developing specific TaqMan ddPCR assays allowing to quantify the frequency of the three *Ae. albopictus Kdr* mutations, *V1016G, I1532T* and *F1534C*, from pools of mosquitoes. The specificity of these assays was validated by using specimens of known genotypes and synthetic DNA constructs for haplotype combinations not yet observed in the field. The accuracy of each assay was validated by comparing *Kdr* frequencies measured from pooled mosquito heads to those obtained from the individualized body of the same individuals. The applicability of these assays for resistance monitoring was then assessed by conducting a pilot surveillance study in mainland France integrating deltamethrin bioassays, ddPCR *Kdr* mutation tracking and the assessment of deltamethrin pressure associated with vector control interventions. The potential of the ddPCR approach for improving the surveillance of *Kdr* resistance alleles at a country-wide scale is then discussed in terms of specificity, sensitivity, accuracy, accessibility, throughput and cost-efficiency.

## Materials and methods

### Mosquito lines

Two reference strains susceptible to deltamethrin and *Kdr*-negative were used as controls for both bioassays and molecular analyses: the SPAM strain, sampled in 2007 in the South-East of France [30], and the SRUN strain, sampled in La Réunion island and maintained in insectaries for more than 20 years [31].

A composite *Ae. albopictus* mosquito line originating from Italy and carrying both *V1016G* and *I1532T Kdr* mutations was used for the development of the ddPCR assays. This composite population was created from various field populations sampled in 2018 and was then subjected to deltamethrin selection at the adult stage for more than 20 generations, in order to increase the frequency of resistance alleles.

Individual mosquito samples collected in a Greek *Ae. albopictus* population in 2023 and carrying the *F1534C Kdr* mutation were also used in the development of the *F1534C* ddPCR assay [23].

### Field mosquito populations

Field sampling was performed over three years, from 2020 to 2022, between April and September by local mosquito control operators (*i*.*e*. EID Rhône-Alpes, EID Méditerranée and Altopictus). Sampling efforts were focused on southern and eastern France because *Ae. albopictus*’ implantation in the country happened mainly through the Italian border [32]. For each population, eggs collection was performed using at least five ovitraps spread over a 500m area to ensure sufficient egg yield and genetic diversity. Eggs were then brought back to the laboratory for rearing under controlled conditions (26±2°C, 80±10% relative humidity and a 14h/10h day/night photoperiod), and adult females were then used for bioassays and molecular analyses. Some samples collected in 2023 and 2024 by the EID-Méditerranée, and in 2025 from the Principality of Monaco were used for molecular analyses only (Table 1).

**Table 1.** Field populations used for deltamethrin bioassays and ddPCR *Kdr* assays.

| Population (locality) | Sample code | Administrative region | 2020 |  | 2021 |  | 2022 |  | 2023-2025 |
| --- | --- | --- | --- | --- | --- | --- | --- | --- | --- |
|  |  |  | Bioassays | ddPCR | Bioassays | ddPCR | Bioassays | ddPCR |  |
| Aiton | AIT | Auvergne-Rhône-Alpes | • | • | • | • | • | • |  |
| Aix-les-Bains | AIX | Auvergne-Rhône-Alpes | • | • |  |  |  |  |  |
| Bourg-en-Bresse | BEB | Auvergne-Rhône-Alpes |  |  |  | • | • | • |  |
| Chambéry | CHY | Auvergne-Rhône-Alpes |  |  |  | • | • | • |  |
| Décines-Charpieu | DEC | Auvergne-Rhône-Alpes |  |  |  |  | • | • |  |
| Saint-Martin-d'Hères | SMH | Auvergne-Rhône-Alpes | • | • | • | • | • | • |  |
| Villeurbanne | VIL | Auvergne-Rhône-Alpes |  | • | • | • | • | • |  |
| Bischheim | BIS | Grand Est | • | • | • | • | • | • |  |
| Bergerac | BER | Nouvelle-Aquitaine |  |  |  |  | • | • |  |
| Pau | PAU | Nouvelle-Aquitaine |  |  |  |  | • | • |  |
| Saint-Médard-en-Jalles | SME | Nouvelle-Aquitaine |  |  |  |  | • | • |  |
| Montpellier | MTP | Occitanie | • |  |  |  |  |  |  |
| Montpellier - La Martelle | MTPM | Occitanie |  |  |  |  | • | • |  |
| Montpellier - Mas Drevon | MTPD | Occitanie |  |  |  |  | • | • |  |
| Montpellier - Nord | MTPN | Occitanie |  |  |  |  | • | • |  |
| Muret | MUR | Occitanie |  |  |  |  | • | • |  |
| Nîmes | NIM | Occitanie |  |  |  |  | • | • |  |
| Pérols | PER | Occitanie |  |  |  |  | • | • |  |
| Rivesaltes | RIV | Occitanie |  |  |  |  | • | • |  |
| Saint-Orens | STO | Occitanie |  |  |  |  | • | • |  |
| Saussan | SAU | Occitanie |  |  |  |  | • | • |  |
| Cannes - Harbor | CANH | Provence-Alpes-Côte-d'Azur |  |  |  |  |  |  | • |
| Marseille - Airport | MARA | Provence-Alpes-Côte-d'Azur |  |  |  |  |  |  | • |
| Marseille - Harbor | MARH | Provence-Alpes-Côte-d'Azur |  |  |  |  |  |  | • |
| Nice - City center | NIC | Provence-Alpes-Côte-d'Azur | • |  |  |  |  |  |  |
| Nice - Airport | NICA | Provence-Alpes-Côte-d'Azur |  |  |  |  |  |  | • |
| Toulon - Harbor | TOUH | Provence-Alpes-Côte-d'Azur |  |  |  |  |  |  | • |
| Monaco | MON | Principality of Monaco |  |  |  |  |  |  | • |

In order to confirm the association of the *V1016G* and *I1532T Kdr* mutations with phenotypical resistance, females from four geographically close populations from the area of Montpellier (MTPD, MTPM, MTPN and SAU) were pooled according to their survival status to deltamethrin during bioassays, allowing the comparison of *Kdr* frequencies between dead and surviving mosquitoes. This resulted in a ‘MTP-dead’ pool with 100 mosquitoes, and a ‘MTP-survivor’ pool with 33 mosquitoes, which were both used for molecular analyses.

### Resistance phenotyping

Deltamethrin resistance was monitored using test tubes bioassays on F1 females from field populations according to the WHO testing protocol [33]. 0.03% deltamethrin-impregnated papers were prepared according to WHO recommendations [34] with the exception that a different silicone oil was used (silicone oil AP 150 Wacker, Sigma-Aldrich, USA). These in-house deltamethrin-impregnated papers induced a mortality *>*80% in both reference susceptible lines over multiple experiments. Bioassays consisted in exposing lots of twenty 3-5 days old non-blood-fed female mosquitoes to 0.03% deltamethrin, for an hour. Mosquitoes were then transferred to insecticide-free test tubes and allowed to recover for 24h with 10% honey solution before mortality was recorded.

The SPAM and SRUN reference strains were used to normalize bioassay data across distinct testing experiments. First, the mean mortality of SPAM and SRUN lines were computed for each testing experiment (*m*_*i*_) together with their global mean across all testing experiments (*M*). A normalization ratio was then calculated for each experiment by dividing the global average *M* by the local average *m*_*i*_. Normalized mortality data were obtained by multiplying raw mortality data from each testing experiment by its respective normalization ratio. Normalized mortality data exceeding 100% were fixed to 100%. As mortality data obtained with our home-made 0.03% deltamethrin papers on the two reference susceptible lines were lower than the expected 100% WHO diagnostic threshold (likely due to the use of a different silicone oil), the susceptibility status of each field population was inferred by comparison with the two reference susceptible lines. Populations with a normalized mortality higher or equal to the lowest value obtained for the reference susceptible lines across all experiments (SRUN 2020, 85%) were considered ‘susceptible’. Populations with a normalized mortality <85% and *>*73.5% (*i*.*e*. the lower bound of the SRUN 2020 95% confidence interval) were considered as ‘possibly resistant’. Populations with normalized mortalities lower than 73.5% were considered ‘resistant’.

### ddPCR assays design

Primers and probes targeting the three *Kdr* mutations *V1016G, I1532T* and *F1534C* carried by the VGSC gene (AALBF5 accession number LOC109421922) were designed in order to avoid polymorphisms that may impair the specific binding of PCR amplification primers and probes based on available pool-seq data obtained from composite populations from various continents (Europe, South-East Asia, South-West Indian ocean and Africa). Such unwanted polymorphisms were manually screened using Strand NGS 3.4 genome viewer (Strand Life Sciences Pvt. Ltd., Bangalore, India). PCR amplification primes and TaqMan probes were then manually designed and their specificity was verified using Primer-BLAST [35] against the AALBF3 and AALBF5 reference assemblies. See S1 Appendix and S2 Appendix for primer and probe sequences and S1 Figure for genomic context.

Given the genomic proximity of the *Kdr* 1532 and 1534 loci, and the probability that they could at some point co-ocur on the same chromosome, a specific multi-probe ddPCR TaqMan assay was designed to detect each mutation regardless of the mutation present at the other locus. This design involves two PCR amplification primers and six probes which are selectively marked depending on which mutation is targeted (see S2 Appendix for primer and probe sequences and S1 Figure for genomic context).

### DNA extractions

#### Technical validation

The ddPCR *V1016G* and *I1532T* assays were validated by comparing the *Kdr* frequencies measured from pooled mosquito heads to those computed from the corresponding individual bodies. For this, thirty 3-5 days old non-blood-fed females of the Italian composite line known to carry both mutations were used. Mosquitoes were decapitated and the heads were gathered together, while bodies remained individualized. Genomic DNA from individual bodies or pooled heads was extracted using the CTAB-chloroform protocol [36], with a 200 µL working volume. Genomic DNA was eluted in 20 µL nuclease-free water, and was quantified using the Qubit HS assay (Thermo Fisher Scientific, USA). Genomic DNA was then diluted down to 2 ng/µL for ddPCR. As one individual body DNA extraction failed, *Kdr* frequencies obtained from 29 body extracts were compared to the frequency obtained from a pool of 30 heads.

This validation protocol could not be replicated for the *F1534C* mutation for lack of whole mosquitoes. Instead, the ddPCR assay was first tested on gDNA samples obtained from individual mosquitoes collected in Greece and carrying this mutation. Given that the *I1532T* and *F1534C* mutations have scarcely been observed together [37], and considering the complexity of the duplex *I1532T* /*F1534C* assay design, synthetic DNA fragments carrying none, either or both mutations on the same strand were synthetized from Genewiz (Germany) to validate the specificity of the duplex probes. In order to obtain a realistic concentration of the target DNA fragment within the template DNA matrix, synthetic DNA were first diluted to 5*×*10^−7^ ng/µL and then mixed to a 1:1 volume with 10 ng/µL plant DNA (*Primula pedemontana*), resulting in a 1:20 million matrix-to-target ratio. This ratio is comparable to the genome-to-target ratio expected in *Ae. albopictus* (186 bp target within a 1.3 Gb genome). Synthetic DNA sequences are presented in S3 Appendix.

#### Field mosquito populations

For each population, gDNA was extracted using the CTAB method from pools of 3-5 days-old non-blood-fed F1 females with sample sizes ranging from 12 to 100 across populations. For each sample the working volume was adapted to the sample size in each batch (V = 200 µL if *n* ≤ 15 and V = 15 *× n* µL if n *>*15, where n is the number of mosquitoes). Pools of mosquitoes were ground using a Retsch MM400 bead grinder (Verder Scientific, Germany) set to 30Hz, during 5 *×* 40 seconds to ensure homogenous grinding. DNA was eluted in varying volumes of nuclease-free water (20 µL up to *n* = 10 mosquitoes, 5 *× n* µL otherwise), quantified using the Qubit BR assay (Thermo Fisher Scientific, USA), then diluted down to 10 ng/µL for ddPCR.

### ddPCR assays

For both *V1016G* and *I1532T* /*F1534C* assays, ddPCR tests were performed on a Bio-Rad QX200 ddPCR system with all reagents obtained from Bio-Rad (USA) and according to the manufacturer’s instructions [38]. For each sample, a ddPCR mix containing primers and TaqMan probes, PCR reagents, the gDNA template and the XhoI restriction enzyme was first made as described in S1 Appendix and S2 Appendix. This mix was then incubated at 37°C for 15 minutes to allow enzymatic gDNA fragmentation. 20 µL of the ddPCR mix and 70 µL of Droplet Generation Oil for Probes (Bio-Rad) were then charged into a DG8 cartridge (Bio-Rad) for droplet generation. Once droplets were generated, 40 µL were transferred to a 96-well plate for amplification on a Bio-Rad CFX96 PCR system (see S1 Appendix and S2 Appendix for conditions). Droplets reading was performed on both FAM and HEX channels using a QX200 Droplet Reader. Droplet assignment to the four categories (HEX-/FAM-, HEX+/FAM-, HEX-/FAM+, and HEX+/FAM+) was performed automatically using the Quantasoft software (Bio-Rad). *Kdr* allele frequencies relative quantities and confidence intervals were then computed by fitting the fraction of negative and positive droplets for each category to a Poisson law as described in the ddPCR Applications Guide [38].

### Linking vector control activities, resistance levels and *Kdr* mutations frequencies

Insecticide selection pressure resulting from vector control interventions was estimated based on the cumulated number of deltamethrin spraying interventions performed by the three operators (EID Méditerranée, EID Rhône-Alpes and Altopictus, data extracted from the national registry for the prevention of vector-borne diseases (SI-LAV)) from June 2020 to December 2024 within their respective area of activity. Adulticid sprayings are realized in 150 to 300 meters radius areas. From this 4-year dataset, the number of sprayings within a 10 km radius from each sampling site was computed, and was used as a proxy of the local insecticide pressure intensity. A 10 km distance was chosen as an estimate of an *Ae. albopictus* population’s dispersion range over a few years (individual maximum flight distance : 676 ± 458 m [39], 5-17 generations per year [40]).

### Statistical analyses

The association between *Kdr* mutation frequencies and deltamethrin resistance in the ‘MTP-survivor’ and ‘MTP-dead’ pools was assessed by performing a Pearson’s chi-square test on droplet counts (`stats` package version 4.5.1). To study the link between vector control activities, *Kdr* mutations frequencies and deltamethrin resistance levels, Pearson’s correlation coefficients were computed between the number of sprayings, the normalized bioassay mortality, and *Kdr* mutation frequencies across all sampling sites (`stats` package version 4.5.1). All analyses were performed in R version 4.5.1 [41].

## Results

### Deltamethrin resistance monitoring by bioassays

A total of 29 field populations sampled from 2020 to 2022 were tested for deltamethrin resistance using diagnostic bioassays as compared to two reference lines (Fig 1 and S1 File). Overall, deltamethrin normalized mortality was relatively high across all populations. According to our resistance thresholds (see Methods), 19 populations (65.5%) were considered susceptible, 10 (34.5%) were considered possibly resistant, and none were considered resistant. Populations considered ‘possibly resistant’ were mostly located in the South of France. The northernmost population, Bischheim (BIS, Grand Est region) was also considered ‘possibly resistant’ in 2021, although the 95% confidence interval was wide.

**Figure 1.**
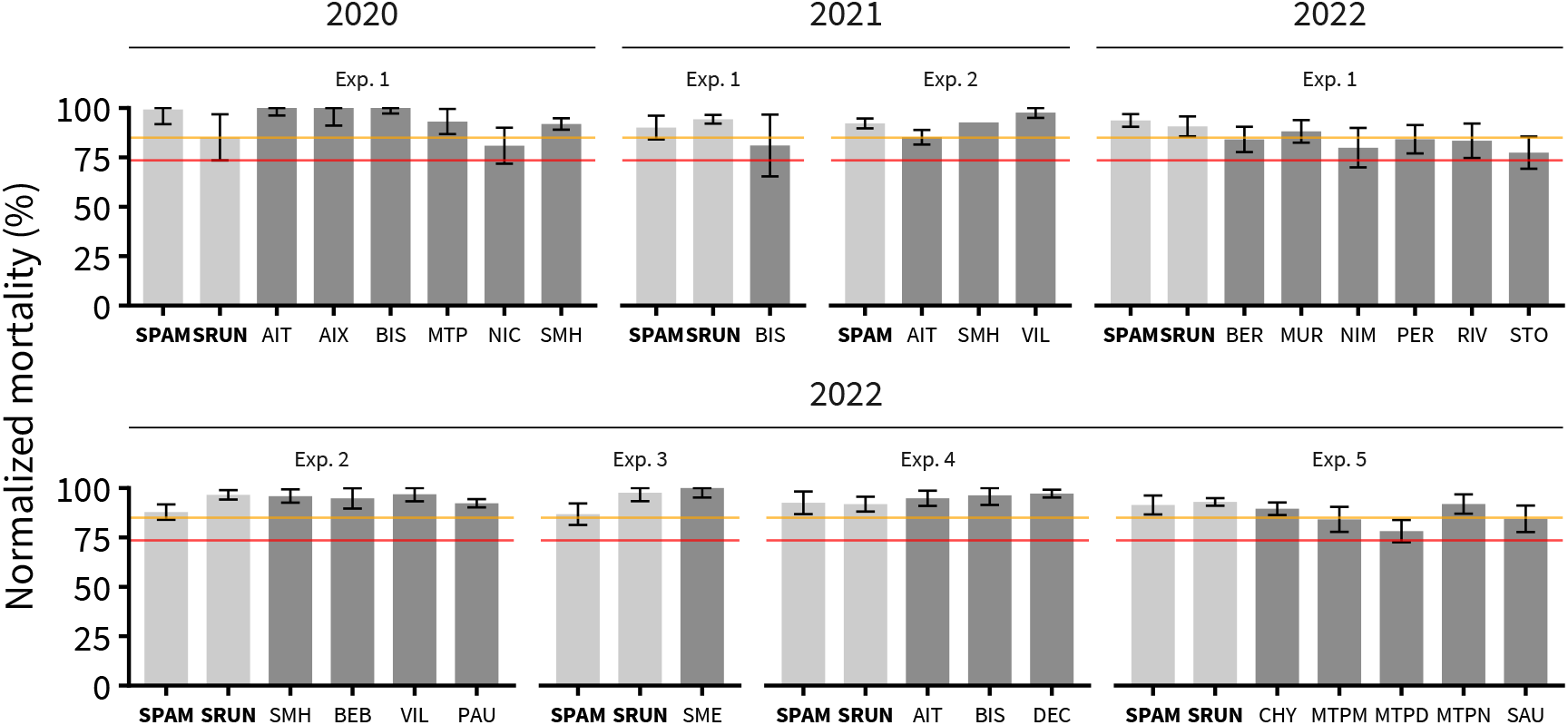
Deltamethrin susceptibility of field *Ae. albopictus* populations. A total of 29 populations sampled between 2020 and 2022 were tested by 0.03% deltamethrin diagnostic bioassays on adult females compared to the two susceptible reference lines SPAM and SRUN (light grey and bold font). Normalized mortality data are shown for each population ±95% confidence intervals. The orange and red lines show the lowest normalized mortality value and the lower bound of its 95% confidence interval obtained from the reference susceptible lines across all testing experiments, respectively.

### ddPCR *Kdr* assays validation

#### *V1016G* and *I1532T* assays

The ddPCR assays performed on individual bodies and pooled heads obtained from adult females of the Italian resistant line confirmed the presence of both *V1016G* and *I1532T Kdr* mutation at medium frequencies (~ 42% and ~ 56% respectively, Fig 2). The frequency of the *1016G* allele inferred from the genotyping of individual body extracts (1 body extraction failed) was very similar to the frequency directly measured from pooled heads (41.4% and 43.2% ± 0.9% respectively). The same trend was observed for the *I1532T* assay with similar frequencies obtained from individual bodies and pooled heads (56.9% and 56.6% ± 0.8% respectively). Overall, this confirmed the ability of the ddPCR to accurately estimate *Kdr* mutation frequencies from pools of mosquitoes. The slight differences observed between *Kdr* frequencies obtained from individual bodies and pooled heads for both assays were likely due to the fact that 29 bodies (one extraction failed) were compared to a single DNA extract obtained from 30 pooled heads.

**Figure 2.**
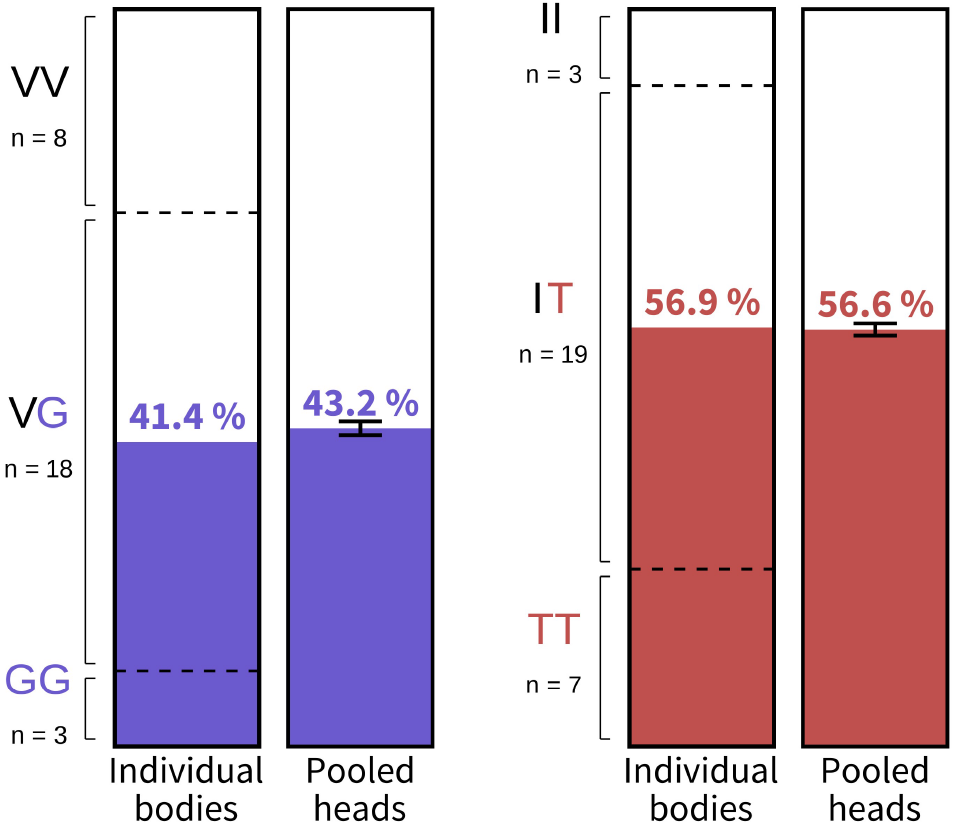
Comparison of the *V1016G* (A) and *I1532T* (B) *Kdr* mutation frequencies obtained by ddPCR assays from individual bodies and pooled heads. DNA extracts were obtained from the same specimens of the Italian resistant line. Genotype counts are shown on the left for individual bodies. Frequencies obtained from pooled heads are shown ± 95% CI computed from the Poisson law.

#### *I1532T* /*F1534C* duplex assay

In order to consider all haplotypes that may occur across the two loci, the specificity of the *I1532T* /*F1534C* duplex ddPCR assay was first tested using four synthetic DNA fragments representing the different haplotypes (I/F, I/C, T/F, T/C) diluted within an exogenous DNA matrix. Overall, the assay showed a good specificity for detecting both mutations within a single reaction, with measured frequencies near 0% and 100% in absence and presence of the resistant alleles respectively (Table 2). The only haplotype showing imperfect specifity was the *1532I* /*1534C* combination, where a minor non-specific signal was observed (1.1% *1532T* and 2.5% *1534F*) though 95% confidence intervals still overlapped 0%. When using the duplex ddPCR assay on DNA samples obtained from single Greek *1532I* /*1534C* mosquitoes, the measured *1532T* and *1534F* frequencies remained as low as 0.09% ± 0.05% and 0.0% ± 0.1% respectively, suggesting that the non-specific effect is negligible when using a realistic mosquito gDNA extract.

**Table 2.** Allelic frequencies obtained from the duplex *I1532T* /*F1534C* assay on the four synthetic DNA for each locus.

| Target allele |  | *1532T* |  | *1534C* |  |
| --- | --- | --- | --- | --- | --- |
|  |  | Freq. (%) | 95% IC | Freq. (%) | 95% IC |
| Haplotype | *1532I/1534F* | 0.0 | - | 0.0 | - |
|  | *1532T/1534F* | 100.0 | 97.8 - 100.0 | 0.0 | - |
|  | *1532I/1534C* | 1.1 | 0.0 - 3.8 | 97.5 | 93.8 - 100.0 |
|  | *1532T/1534C* | 100.0 | 98.4 - 100.0 | 100.0 | 98.3 - 100.0 |

Upper confidence interval bounds have been truncated down to 100% when they were higher.

### Implementation of the ddPCR *Kdr* assays on field populations

ddPCR *Kdr* assays were implemented on the 30 field populations from mainland France and Monaco for which DNA extracts were available (for sample information, see S1 File). The *1534C Kdr* allele was not detected while *V1016G* and *I1532T* alleles were detected at low frequencies (Fig 3 and S1 File). The *V1016G* allele was detected in only 5 field populations out of 30 (16.6%), with frequencies ranging from 0.52% (TOUH) to 3.88% (MON). These 5 populations were located in the South of France and were also carrying the *I1532T* mutation. The *1532T* allele was detected in 19 populations our of 30 (63.3%), with frequencies ranging from 0.35% (NIM) to 12.93% (BER).

**Figure 3.**
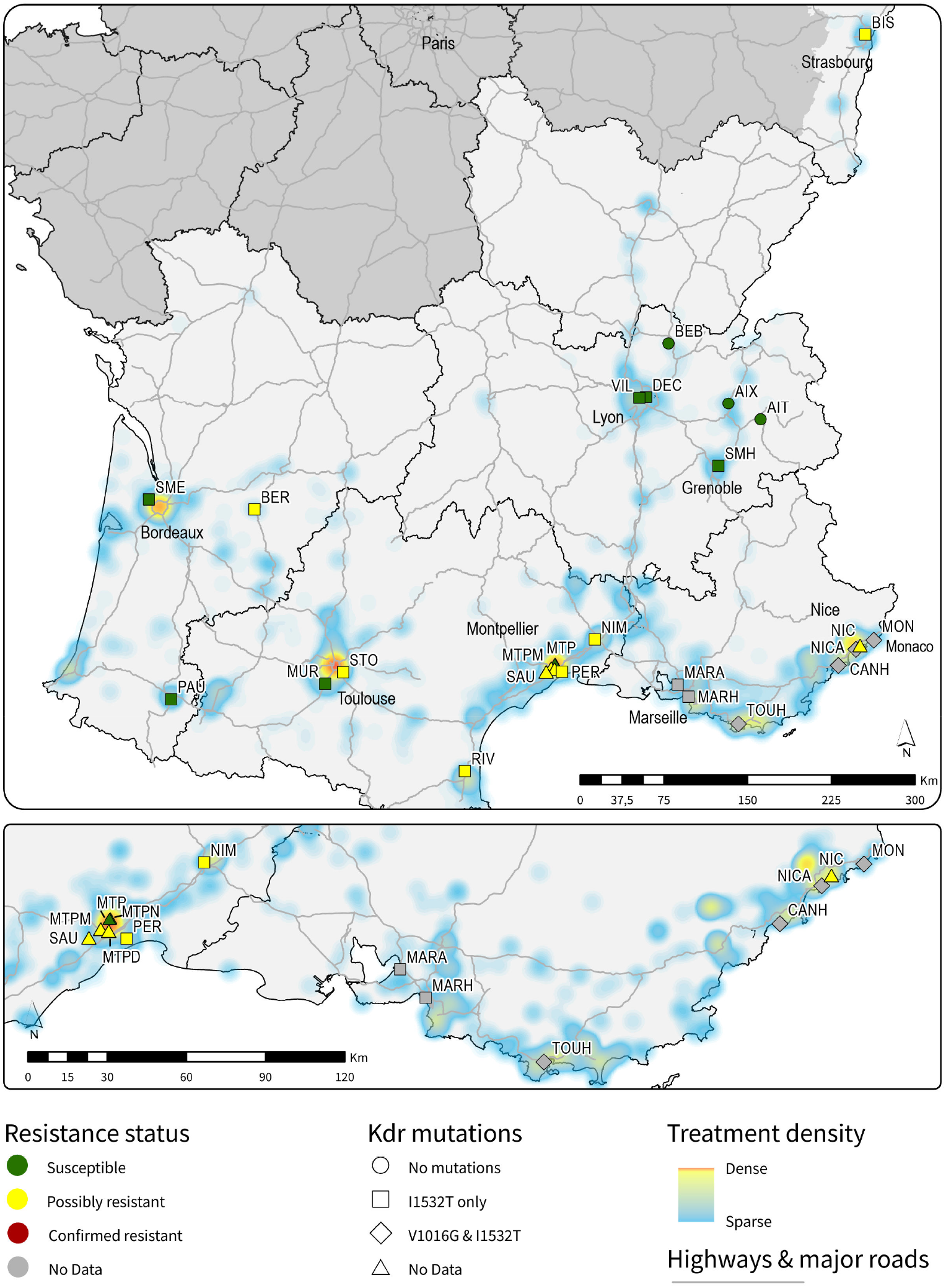
Spatial representation of deltamethrin resistance and *Kdr* mutations occurrence in regards to deltamethrin interventions in mainland France and Monaco. Deltamethrin interventions data only included curative vector control interventions performed by accredited mosquito control operators in the study area from June 2020 and December 2024. Deltamethrin resistance status refers to those inferred from normalized mortality rates generated from bioassays with 0.03% deltamethrin, and *1016G* and *1532T* resistance alleles occurence is indicated when applicable. Administrative regions are delimited by black lines and major traffic roads (highways and main national roads) are shown in light grey. Regions in dark grey were not covered by the present study. For the sake of clarity, when a sampling site has been used over several years, only the lowest mortality and the highest *Kdr* mutation frequencies are shown.

The genotype/phenotype association study conducted on dead and survivors from the four populations close to Montpellier in 2022 (MTPD, MTPM, MTPN and SAU) supported the association of both *1016G* and *1532T* alleles with resistance. The ‘MTP-survivor’ pool, showed a higher frequency of both *V1016G* and *I1532T* alleles (2.69% ± 0.41% and 3.86% ± 0.60% respectively) compared to the ‘MTP-dead’ pool (0.00% and 0.95% ± 0.25% respectively, Pearson’s chi-square test p-value ≤ 0.001 for both alleles).

### Relationship between deltamethrin treatments, resistance levels and *Kdr* frequencies

A total of 1800 deltamethrin space spraying interventions were carried out by local mosquito control operators in the context of emergency/curative vector control within the study area between June 2020 and December 2024. 312 spraying interventions were carried out in 2020, 53 in 2021, 277 in 2022, 581 in 2023 and 577 in 2024, showing an increase over the years. Among them, 529 (29.4%) were recorded within 10 km of a sampling site. The number of sprayings per site ranged from 0 (AIT, BEB) to 118 (MTP and MTPN), with an average of 43.5 sprayings per site. More than 80% of spraying interventions occurred within 10 km of a highway, indicating that arboviral cases and subsequent deltamethrin interventions tend to occur close to major communication axes linking important urban areas. Looking for pairwise correlations between the number of deltamethrin interventions within 10 km of sampling sites, normalized mortality from bioassays and *Kdr* mutations frequencies obtained by ddPCR across the 28 informative sites did not evidence any significant correlation between any of the three factors. Given the low frequencies of *Kdr* alleles, correlations were also computed with presence/absence data instead of continuous *Kdr* frequencies. All correlations remain non-significant except for the one observed between the *1532T* allele presence and the number of deltamethrin interventions (p = 0.031). Correlation results for both continuous and binary *Kdr* data are presented in Table 3.

**Table 3.**
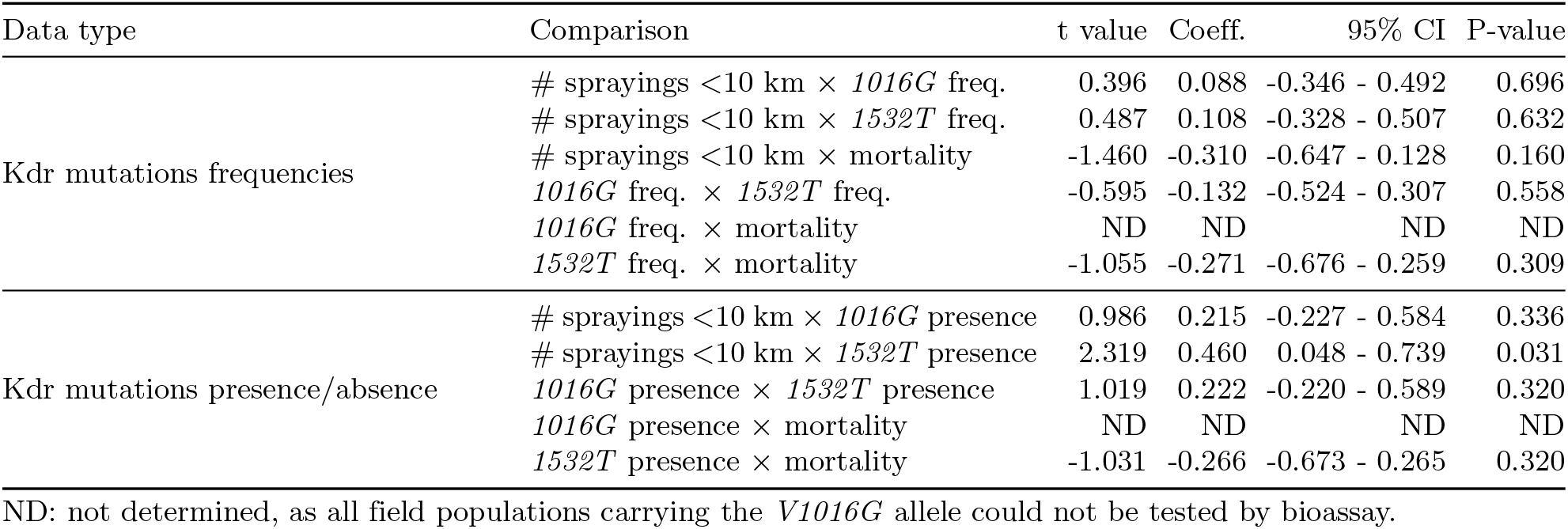
Pearson’s pairwise correlation statistics.

## Discussion

### Novel ddPCR assays for the monitoring of *Kdr* mutations frequencies at high-throughput in *Ae. albopictus*

Due to its high genetic plasticity and adaptation capabilities, and global climate change, *Aedes albopictus*’ distribution range is expanding at a very fast pace [42, 43]. Such expansion led to an increase of arboviral diseases transmission such as dengue and chikungunya in new territories including EU countries [44]. For instance, autochtonous chikungunya cases in France went from 17 cases in 2017 to 809 cases in 2025 [5]. In the absence of efficient vaccinal strategies, vector control remains an immediate and the most efficient way to prevent outbreaks. Though biological control with *Bacillus thuringiensis var. israelensis* (Bti) toxins can efficiently target larvae and alternative control tools are being developed (*e*.*g. Wolbachia*-based arbovirus transmission blocking, sterile or incompatible insects release, mass trapping), and while they are efficient preventative measures, they are not suitable for curative vector control. Pyrethroids are still consequently routinely used to target adults to control arbovirus transmission in high-risk contexts. Considering the increased pyrethroid resistance currently observed in *Ae. albopictus* worldwide [45] and the speed at which *Kdr* mutations spread in populations under selective pressure ([46]), the regular monitoring of major resistance alleles such as *Kdr* mutations in conjunction with bioassays represents a key component of resistance management strategies.

Current PCR-based *Kdr* genotyping assays require the processing of single mosquito specimens from gDNA extractions to PCR and data interpretation [22, 47, 48]. In order to reach a high sensitivity in *Kdr* mutations detection and accurately quantify their frequencies in natural populations, at least 50 mosquitoes per population (2N=100) would be required, making such approach laborious and costly to reach sufficient coverage at a country-wide scale (e.g. ~30 populations making 30 *×* 50 = 1500 gDNA reactions annually).

In this respect, the ddPCR Taqman *Kdr* assays we developed undeniably represent a promising way to improve pyrethroid resistance surveillance.

First, these assays were designed in order to be applicable to various mosquito genetic backgrounds with minimum interference of amplification primers and TaqMan probes with polymorphisms adjacent to the targeted mutations. Although such polyvalence still needs to be experimentally validated on mosquitoes from various continents, ongoing studies confirmed that these assays are readily applicable to *Ae. albopictus* populations from various EU countries (*i*.*e*. mainland France, Italy, Switzerland and Greece).

Second, these assays show a good specificity for the detection of the three major *Kdr* alleles that have been associated with resistance in *Ae. albopictus* (*i*.*e. 1016G, 1532T* and *1534C*). The three *V1016G, I1532T* and *F1534C* single-plex assays were highly specific to the targeted allele, while the duplex *I1532T* /*F1534C* assay only showed a negligible cross-signal between probes allowing to detect both *1532T* and *1534C* alleles even if they are genetically linked (*i*.*e*. located on the same chromosome strand). Though the existence of a double-mutant haplotype *1532T* /*1534C* in wild mosquito populations is still unclear [49], its emergence will not affect the ability of the assay to accurately quantify both mutations. Nevertheless, if only one of either the *I1532T* or *F1534C* mutations is expected to be present in a given population, the test can also be decoupled by using only the relevant TaqMan probes.

Third, by partitioning the gDNA sample in thousands of nanodroplets, the ddPCR technology allows reaching a very high sensitivity and accuracy in quantifying *Kdr* mutations’ frequencies provided that the number of specimens constituting the DNA pool is sufficient. For instance, both *V1016G* and *I1532T* assays yielded very close allelic frequencies when applied to pooled heads and individual bodies from the same mosquitoes, indicating that both tests can be reliably used on pooled specimens. The use of these assays on field samples made from up to 100 specimens per pool was also successful and allowed detecting resistance alleles at very low frequencies. Considering an average of 3000 positive droplets per reaction and a threshold of 5 positive droplets per channel, the assay would be sufficient to detect a single heterozygote mutant within a pool of 300 specimens (600N chromosomes), lowering down its sensitivity to a 0.16% *Kdr* mutation frequency.

Fourth, considering that *Kdr* mutations frequencies are sufficient to track resistance alleles, the ability of these assays to be applied on pooled specimens allows a significant increase of throughput by reducing time and efforts associated with DNA extractions, PCR reactions and data analysis. In our case our laboratory setting allowed us to extract up to 48 pools of mosquitoes in a single day, making the whole process from sample reception to results less than two days (*i*.*e*.DNA extraction, quality check and dilution; ddPCR assay and result interpretation). However, as the PCR step can handle up to 96 samples, further optimizing DNA extraction throughput will further increase the sample flow rate.

Altogether, we believe these novel TaqMan ddPCR *Kdr* assays meet the requirement to enhance the routine surveillance of spatiotemporal dynamic of pyrethroid resistance alleles at a country scale.

Finally, the ability of these ddPCR TaqMan assays to be performed on pools of mosquitoes allows a significant reduction of time-to-result and costs as compared to other *Kdr* genotyping assays (Table 4 and S1 Table). In addition to reducing human costs through the fast processing of a limited number of samples, a significant cost reduction is also achieved on reagents and consumables. Based on our estimation, the cost gain for processing 20 populations of 100 mosquitoes can reach 97% and 95% compared to common genotyping approaches (440€ for ddPCR vs 15200€ and 9010€ for Sanger sequencing and qPCR TaqMan respectively). A similar cost reduction can be expected compared to genotyping by multiplexed mass sequencing though the cost of this approach may be lowered by the concomitant sequencing of multiple *Kdr* loci as sequencing costs are negligible. However, such cost gain should be put in perspective of the ddPCR equipment profitability (~50K€ investment for a dual channel ddPCR system). In terms of time-to-result, the ddPCR assay largely outperforms other approaches with *Kdr* allele frequencies obtained for the 20 populations in only 2 working days. Other approaches show longer time-to-result due to the high number of samples to proceed, delays associated with sequencing subcontracting (Sanger sequencing or Mass sequencing) and bioinformatic analyses (Mass sequencing). Considering the above, the use of ddPCR TaqMan assays on pool of mosquitoes appears as a high-throughput and cost-saving approach for the routine surveillance of *Kdr* mutations in *Ae. albopictus* at a country scale.

**Table 4.** Cost estimates of traditional genotyping methods compared to the TaqMan ddPCR pooled approach.

| Approach | Reagent costs (€) | Personnel costs (€) | Total cost (€) | Time-to-results | Data type |
| --- | --- | --- | --- | --- | --- |
| Individual genotyping by Sanger sequencing | 12200 | 3010 | 15210 | >1 month | Genotypes |
| Individual genotyping by TaqMan qPCR | 6000 | 3010 | 9010 | ~20 days | Genotypes |
| Individual genotyping by mass multiplex short read sequencing | 9400 | 3610 | 12810 | ~2 months | Genotypes |
| TaqMan ddPCR on pools | 140 | 300 | 440 | 2 days | Allele frequencies |

Cost and time-to-results were estimated for the screening of 1 *Kdr* mutation from 20 *Ae. albopictus* populations each comprised of 100 specimens. Costs are expressed in euros and rounded to the nearest ten. Time-to-results for sequencing methods has been estimated including subcontracting processing time. Detailed calculations are provided in S1 Table.

### Outcome of the pilot study in France

Through a partnership with local mosquito control operators, a pilot resistance surveillance study integrating deltamethrin bioassays, the ddPCR screening of *Kdr* mutations circulating in mainland France (*V1016G* and *I1532T*) and an assessment of pyrethroid pressure related to vector control interventions was conducted.

Compiling vector control intervention records from June 2020 to December 2024 confirmed the increasing pyrethroid pressure associated with public-health interventions in the southern part of France. Considering that pyrethroids are also used by private pest control companies, at home and for pest control in agriculture, such increase in selection pressure is likely underestimated. As expected, areas showing the highest pyrethroid pressure over the last years were located in the southern part of France near densely populated areas (Nice, Toulon, Marseille, Montpellier, Toulouse and Bordeaux) where high *Ae. albopictus* density and arbovirus transmission cases were reported.

Phenotypical resistance data obtained from bioassays suggest that deltamethrin resistance prevalence remains moderate in French *Ae. albopictus* populations and that vector control interventions are still effective for controlling arbovirus transmission. The overall resistance alleles distribution seems consistent with their Italian origin [17] with most populations exhibiting resistance signals being located in an east-west transect from the Italian border. Although a significant but weak correlation was observed between deltamethrin interventions and the presence of the *Kdr 1532T*, this result must be interpreted with caution as it likely reflects spatial autocorrelation rather than a causal link between the insecticide selection pressure resulting from local mosquito control interventions and the *1532T* allele. Indeed, deltamethrin interventions are mainly concentrated in the south-east part of France, which is associated with an important gene flow from Italian populations where *Kdr* mutations are circulating at higher frequencies. While no other significant correlation was observed between vector control interventions and deltamethrin resistance, such result should be taken with caution considering the limited accuracy of bioassays and the limited number of populations tested from the South-East because of sampling constraints.

The implementation of ddPCR TaqMan assays to monitor the frequency of *V1016G* and *I1532T Kdr* mutations confirmed their circulation in mainland France. The *1016G* resistant allele seems to be circumscribed to the South-East region, which is consistent with its supposed point of entry, through the Italian border. This pattern is consistent with previous studies using either individual PCR genotyping or mass sequencing [18, 19]. The *1532T* resistant allele seems to be present at a higher frequency and more widely distributed as it was detected in most regions. Its detection near Lyon and up to Strasbourg is in line with prior findings [24]. Such higher prevalence of the *1532T* resistant allele may also reflect its lower fitness cost under moderate insecticide pressure. In this regard, although a significant frequency was observed for both *1016G* and *1532T* alleles in the deltamethrin-selected Italian line, no double homozygote was observed, suggesting that these two alleles might be carried by distinct haplotypes and that their cumulated fitness cost may limit their genotypic association. Though a significant enrichment of both resistant alleles was observed in deltamethrin survivors from the Montpellier area, the absence of a strong correlation between vector control interventions and both phenotypic and molecular resistance data suggest that until then, the restricted deltamethrin spraying policy applied by vector control operators was successful at limiting the selection of resistance alleles. However, considering the increase of arboviral transmission cases and subsequent deltamethrin vector control interventions in the last few years, in addition to nuisance control, this situation may rapidly evolve and calls for an increased monitoring of resistance at the country scale. In this regard, the novel TaqMan ddPCR assays we developed could become be a key component of resistance surveillance programs.

### Towards an integrated pyrethroid resistance surveillance framework in mainland France and beyond

Considering the establishment of *Ae. albopictus* in mainland France, the increased usage of deltamethrin to prevent arbovirus outbreaks and the presence of known *Kdr* resistance alleles, there is an urgent need to establish a sustainable resistance surveillance program for the upcoming decade until alternative arbovirus control strategies are widely implemented. Ideally, such surveillance program should involve a partnership between all public health operators operating through the country and the research community, and be supported by public health agencies such as the French agency for food and health security (ANSES) and the Health ministry department (DGS).

In terms of framework, such a resistance surveillance program should generate annual or biennial data, extend over a sufficient duration (*i*.*e*. ~10 years) and adequately cover the country’s territory to provide a meaningful picture of the spatio-temporal dynamics of pyrethroid resistance in *Ae. albopictus* populations. Such program will highly benefit from the integration of phenotypic resistance data obtained from standardized bioassays, *Kdr* mutations frequencies and the assessment of pyrethroid pressures. In terms of implementation, mosquito control operators are highly experienced with field collections and could provide the biological material (*i*.*e*. mosquito eggs) necessary for performing bioassays and molecular work, as part of their existing field activities. From this material and in order to limit personnel costs, deltamethrin bioassays could be performed on a well-defined set of sentinel populations (*i*.*e*. ~10-15 established populations spread over the country and matching main communications routes and entry points). The monitoring of *Kdr* mutations’ frequencies could be performed from a larger set of populations (*i*.*e*. up to 50 populations) for a moderate cost using ddPCR assays. If needed, additional individual mosquito specimens may also be used to investigate *Kdr* genotype frequencies for a subset of populations of interest.

As shown in the present study, pyrethroid pressure intensity assessment can be readily achieved by compiling deltamethrin intervention records performed by accredited mosquito control operators over a few years and may also be refined by integrating additional data from private pest control companies and agriculture. Finally, such dataset may also be related to operational deltamethrin efficacy assessment from operators and eventually to arboviral transmission rates.

Overall, the implementation of such long-term surveillance program will allow predicting resistance risk and anticipating the deployment of management actions for a limited cost. In the medium term, extending such surveillance framework to other EU countries could also be achievable through partnerships across the EU mosquito control and research community that should be facilitated by existing networks and public health agencies (*i*.*e*.the European Mosquito Control Association, the European Society for Vector Ecology and the European Centre for Disease Prevention and Control).

## Conclusion

The novel ddPCR TaqMan assays developed here offer a great opportunity for the high-throughput monitoring of *Kdr* mutations in *Ae. albopictus* populations at a country scale, which appear as a key component of insecticide resistance surveillance programs. These assays are cost-effective, time-efficient, highly sensitive and specific, and versatile enough to be used both for individual genotyping and large-scale pooled surveillance. The implementation of these tests within a pilot study from French *Ae. albopictus* field populations confirmed the presence of both *V1016G* and *I1532T* mutations at low frequencies with a minor effect on phenotypic resistance. However, considering the current epidemiological context and the increasing use of pyrethroids for vector and nuisance control, the present study calls for improving resistance surveillance at the country scale and beyond.

## Acknowledgments

The authors acknowledge the support of the local mosquito control operators (EID Méditerranée, EID Rhône-Alpes, SLM67 and Altopictus) for their help in field sampling, and providing deltamethrin interventions data with permission from the local health agencies (ARS) and the General Directorate of Health (DGS). Moreover, we thank the ARS Occitanie, ARS Nouvelle-Aquitaine, ARS Provence-Alpes-Côte d’Azur, ARS Rhône-Alpes and ARS Grand-Est and their field agents for their support in sampling field mosquitoes, including by coordinating transportation during routine monitoring missions. We would also like to thank the following individuals for their contribution to the sampling efforts: Christophe Robino (Département des Affaires Sociales et de la Santé), Eric J. Voiglio (Direction de l’action sanitaire), Jérôme Durivaut, Guillaume Groshenry and Christian Lavagna (Centre Scientifique de Monaco) for Monaco, Lionel Chanaud and Yves-Marie Kervella for the EID Méditerranée. Finally, we thank Alessandra della Torre and Paola Serini (Sapienza University of Rome) for their contribution to the sampling and rearing of the Italian composite line.

## Supporting information

**S1 Appendix. Standard operating procedure for the quantification of *Kdr V1016G* allelic frequency using ddPCR**.

**S2 Appendix. Standard operating procedure for the quantification of *Kdr I1532T* and *F1534C* allelic frequencies using ddPCR**.

**S1 Figure. ddPCR TaqMan oligonucleotides positions on the AALBF5 reference genome. S3 Appendix. Synthetic DNA sequences used for ddPCR validation**.

**S1 File. Bioassay mortality, *Kdr* mutations frequency and deltamethrin interventions data of field populations**.

**S1 Table. Cost estimations of traditional genotyping tools compared to the novel ddPCR TaqMan assays**.

