## Supplementary material for "Novel ddPCR diagnostic assays for the high-throughput surveillance of *Kdr* mutations in *Aedes albopictus*": S1 Appendix

### Supplementary file 1: Quantification of kdr V1016G allelic frequency using ddPCR

#### 1 Primers and probes

| Target allele | Oligo name | Oligo sequence (5' to 3') |
|---|---|---|
| 1016V (wild type) + 1016G (mutated) | F_V1016G | ACCGTAGTGATAGGAAATCTAG |
| | R_V1016G | CGCGATCTTGTTCG |
| 1016V (wild type) | 1016V* | [HEX]CCAGG**T**ACTTAACCTTTTCTTAGC[BHQ1] |
| 1016G (mutated) | 1016G* | [6FAM]CCAGG**G**ACTTAACCTTTTCTTA[BHQ1] |

#### 2 ddPCR Mix

| | Initial concentration | Volume per well (µL) |
|---|---|---|
| ddPCR Supermix for Probes (No dUTP) | 2X | 12.50 |
| F_V1016G | 10 µM | 1.00 |
| R_V1016G | 10 µM | 1.00 |
| 1016V* | 10 µM | 0.25 |
| 1016G* | 10 µM | 0.25 |
| Restriction enzyme XhoI | 10 U/µL | 0.25 |
| rCutSmart buffer | 10X | 0.11 |
| gDNA | * | * |
| RNAse-free water | - | q.s. 25 µL |
| | **TOTAL VOLUME** | **25 µL** |

*Target DNA quantity per well is around 10 ng.

#### 3 ddPCR conditions

| Step | Temperature | Duration |
|---|---|---|
| 1. Polymerase activation | 95°C | 10 minutes |
| 2. Denaturation | 95°C | 30 seconds |
| 3. Annealing + Extension | 56°C | 2 minutes |
| 4. Repeat steps 2. and 3. 39 times | | |
| 5. Polymerase inactivation | 4°C | 5 minutes |
| 6. Polymerase inactivation | 90°C | 5 minutes |
| 7. Post-PCR idle time | 4°C | ∞ |

Temperature ramping was set to 2°C/second.

#### 4 Scatterplots

##### 4.1 Scatterplot genotyping

Example scatterplots for the three kdr V1016G genotypes are presented below. In each panel, green dots are droplets containing the wild-type amplicon 1016V only, blue dots the droplets containing the mutated amplicon 1016G only, orange dots the droplets containing both amplicons, and grey dots indicate droplets containing neither. The HEX channel correponds to the wild-type allele 1016V, while the FAM channel correponds to the mutated allele 1016G.

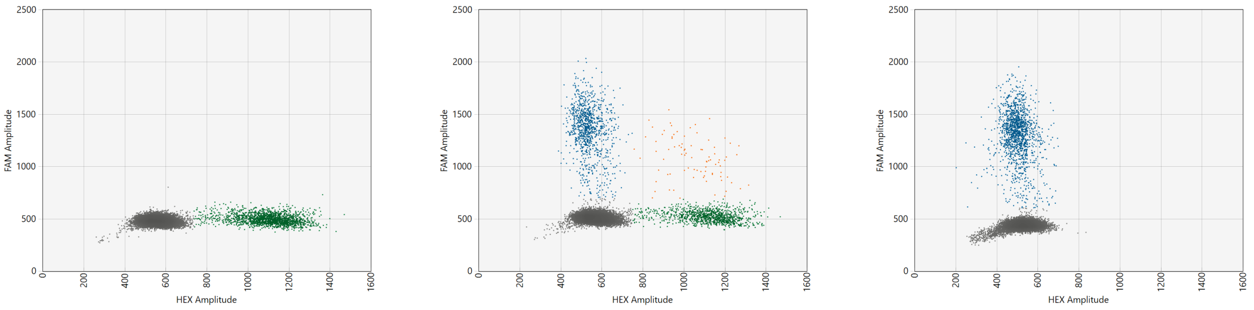


**(a)** 1016V homozygote **(b)** 1016G heterozygote **(c)** 1016G homozygote

#### 4.2 Scatterplot of a pooled sample

Example scatterplot obtained from a pool of 100 mosquitoes of the Italian resistant composite population with the V1016G assay. The estimated frequency of the 1016G allele in this pool is 31.8%.

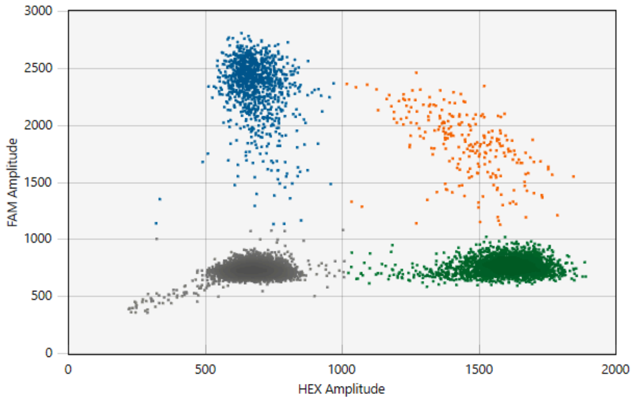


### 5 Estimation of the allelic frequency

ddPCR partitions (droplets) are classified as positive or negative by the reader for each allele, which follow a Poisson distribution. From these, Poisson statistics are then used to estimate the absolute concentration of each allele in the reacion mix, which in turn allow the calculation of the frequency of the mutated allele. Confidence interval bounds are calculated using the Poisson distribution in the same manner.
