## Supplementary material for "Novel ddPCR diagnostic assays for the high-throughput surveillance of *Kdr* mutations in *Aedes albopictus*": S2 Appendix

### Supplementary file 2: Quantification of kdr I1532T/F1534C allelic frequency using ddPCR

#### 1 Primers and probes

| Target haplotype | Target locus | Oligo name | Oligo sequence (3' to 5') |
|---|---|---|---|
| 1532I + 1532T + 1534F + 1534C | 1532 + 1534 | F_1532_1534 | GCGAGACCAACATCTACA |
| 1532I + 1532T + 1534F + 1534C | 1532 + 1534 | R_1532_1534 | AACATTTCCAGCGAGC |
| 1532I + 1534F | 1532 + 1534 | 1532I*_1534F* | [HEX]GTGTTCTTCA**T**CATCT**T**CGG[BHQ1] |
| 1532I + 1534C | 1532 | 1532I*_1534C | [HEX]GTTCTTCA**T**CATCT**G**CGG[BHQ1] |
| 1532T + 1534F | 1532 | 1532T*_1534F | [FAM]TCTTCA**C**CATCT**T**CGGG[BHQ1] |
| 1532T + 1534F | 1534 | 1532T_1534F* | [HEX]TCTTCA**C**CATCT**T**CGGG[BHQ1] |
| 1532I + 1534C | 1534 | 1532I_1534C* | [FAM]GTTCTTCA**T**CATCT**G**CGG[BHQ1] |
| 1532T + 1534C | 1532 + 1534 | 1532T*_1534C* | [FAM]TCTTCA**C**CATCT**G**CGG[BHQ1] |

#### 2 ddPCR Mix

| | Initial concentration | Volume per well (µL) |
|---|---|---|
| ddPCR Supermix for Probes (No dUTP) | 2X | 12.50 |
| F_1532_1534 | 10 µM | 1.00 |
| R_1532_1534 | 10 µM | 1.00 |
| 1532I*_1534F* | 10 µM | 0.25 |
| 1532I*_1534C | 10 µM | 0.25 |
| 1532T*_1534F | 10 µM | 0.25 |
| 1532T_1534F* | 10 µM | 0.25 |
| 1532I_1534C* | 10 µM | 0.25 |
| 1532T*_1534C* | 10 µM | 0.25 |
| Restriction enzyme XhoI | 10 U/µL | 0.25 |
| rCutSmart buffer | 10X | 0.11 |
| gDNA | * | * |
| RNAse-free water | - | q.s. 25 µL |
| | **TOTAL VOLUME** | **25 µL** |

*Target DNA quantity per well is around 10 ng.

#### 3 ddPCR conditions

| Step | Temperature | Duration |
|---|---|---|
| 1. Polymerase activation | 95°C | 10 minutes |
| 2. Denaturation | 95°C | 30 seconds |
| 3. Annealing + Extension | 50°C for I1532T<br>54°C for F1534C | 2 minutes |
| 4. Repeat steps 2. and 3. 39 times | | |
| 5. Polymerase inactivation | 4°C | 5 minutes |
| 6. Polymerase inactivation | 90°C | 5 minutes |
| 7. Post-PCR idle time | 4°C | ∞ |

Temperature ramping was set to 2°C/second.

### 4 Scatterplots

#### 4.1 I1532T assay:

##### 4.1.1 Scatterplot genotyping

Example scatterplots for the three kdr I1532T genotypes are presented below. In each panel, green dots are droplets containing the wild-type amplicon 1532I only, blue dots the droplets containing the mutated amplicon 1532T only, orange dots the droplets containing both amplicons, and grey dots indicate droplets containing neither. The HEX channel correponds to the wild-type allele 1532I, while the FAM channel correponds to the mutated allele 1532T.

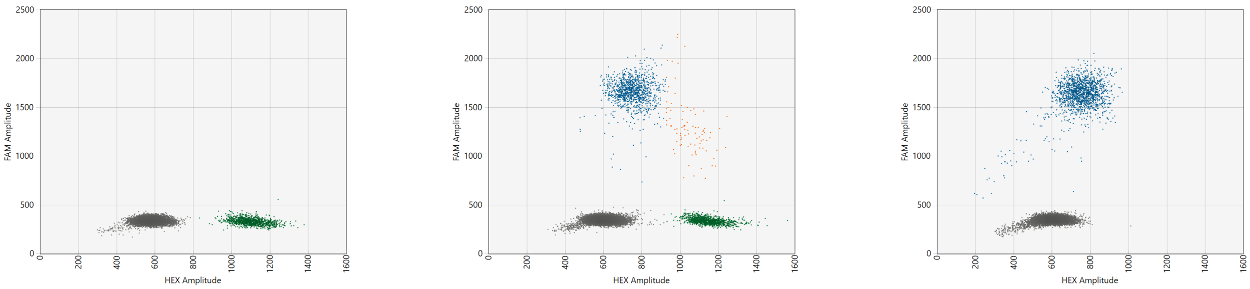


**(a)** 1532I homozygote **(b)** 1532T heterozygote **(c)** 1532T homozygote

##### 4.1.2 Scatterplot of a pooled sample

Example scatterplot obtained from a pool of 100 mosquitoes of the Italian resistant composite population with the I1532T assay. The estimated frequency of the 1532T allele in this pool is 68.4%.

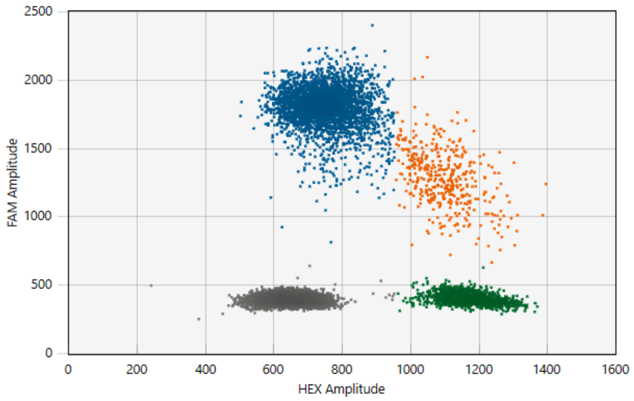

#### 4.2 F1534C assay:

Example scatterplots for the three kdr 1534 genotypes are presented below. In each panel, green dots are droplets containing the wild-type amplicon 1534F only, blue dots the droplets containing the mutated amplicon 1534C only, orange dots the droplets containing both amplicons, and grey dots indicate droplets containing neither. The HEX channel correponds to the wild-type allele 1534F, while the FAM channel correponds to the mutated allele 1534C.

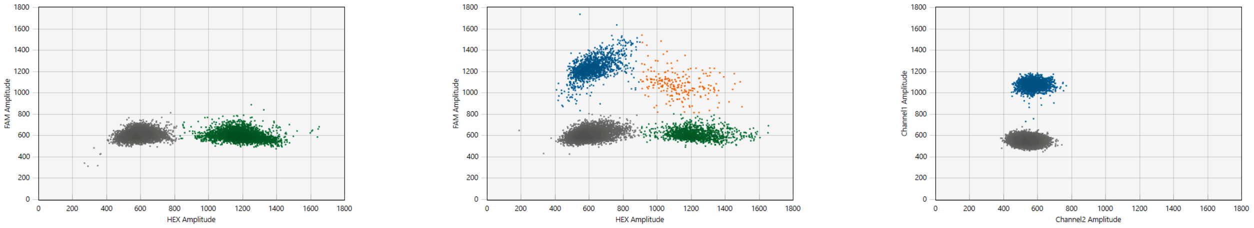


**(a)** 1534F homozygote

**(b)** 1534C heterozygote

**(c)** 1534C homozygote

### 5 Estimation of the allelic frequency

ddPCR partitions (droplets) are classified as positive or negative by the reader for each allele, which follow a Poisson distribution. From these, Poisson statistics are then used to estimate the absolute concentration of each allele in the reacion mix, which in turn allow the calculation of the frequency of the mutated allele. Confidence interval bounds are calculated using the Poisson distribution in the same manner.
