## Supplementary material for "Novel ddPCR diagnostic assays for the high-throughput surveillance of *Kdr* mutations in *Aedes albopictus*": S1 Figure

**Supplementary file 3: ddPCR TaqMan oligonucleotides on the AALBF5 reference genome**

## V1016G :

```
                                                 V1016G
               345229181                       345229043                          345228976
                   v                               v                                  v
WT sequence 3' ACCGTAGTGATAGGAAATCTAG ——————— CCAGGTACTTAACCTTTTCTTAGC ——————— CGAACAAGATCGCG 5'
      F_V1016G ACCGTAGTGATAGGAAATCTAG
        1016V*                                 CCAGGTACTTAACCTTTTCTTAGC
        1016G*                                 CCAGGGACTTAACCTTTTCTTA
      R_V1016G                                                                 CGAACAAGATCGCG
               <———————————————————————————— 208 bp ————————————————————————————>
```

## I1532T :

```
                                                I1532T    F1534C
                   345176281                  345176239  345176233                   345176140
                       v                              v     v                            v
WT sequence 3' GCGAGACCAACATCTACA ——————— GTGTTCTTCATCATCTTCGGG ——————— GCTCGCTGGAAATGTT 5'
   F_1532_1534 GCGAGACCAACATCTACA
 1532I*_1534F*                           GTGTTCTTCATCATCTTCGG
  1532I*_1534C                             GTTCTTCATCATCTGCGG
  1532T*_1534F                               TCTTCACCATCTTCGGG
 1532T*_1534C*                               TCTTCACCATCTGCGG
   R_1532_1534                                                             GCTCGCTGGAAATGTT
               <—————————————————————————— 142 bp ——————————————————————————>
```

## F1534C :

```
                                                I1532T    F1534C
                   345176281                  345176239  345176233                   345176140
                       v                              v     v                            v
WT sequence 3' GCGAGACCAACATCTACA ——————— GTGTTCTTCATCATCTTCGGG ——————— GCTCGCTGGAAATGTT 5'
   F_1532_1534 GCGAGACCAACATCTACA
 1532I*_1534F*                           GTGTTCTTCATCATCTTCGG
  1532I_1534C*                             GTTCTTCATCATCTGCGG
  1532T_1534F*                               TCTTCACCATCTTCGGG
 1532T*_1534C*                               TCTTCACCATCTGCGG
   R_1532_1534                                                             GCTCGCTGGAAATGTT
               <—————————————————————————— 142 bp ——————————————————————————>
```
