## Supplementary material for "Novel ddPCR diagnostic assays for the high-throughput surveillance of *Kdr* mutations in *Aedes albopictus*": S3 Appendix

**Supplementary file 4: Synthetic DNA sequences used for ddPCR validation**

Bold letters represent the positions of the mutations of interest,I1532T and F1534C.

- 1532I/1534F:

  GGAACCAGGTGGGCAAGCAGCCAATTCGCGAGACCAACATCTACATGTACCTCTACTTCGTGTTCTTCA
  **T**CATCT**T**CGGGTCGTTCTTCACCCTTAATCTGTTCATCGGTGTCATCATCGACAACTTCAACGAGCAGA
  AGAAGAAAGCCGGTGGCTCGCTGGAAATGTTCATGACGGAGGATCAGAAAAAGGTTCC

- 1532T/1534F:

  GGAACCAGGTGGGCAAGCAGCCAATTCGCGAGACCAACATCTACATGTACCTCTACTTCGTGTTCTTCA
  **C**CATCT**T**CGGGTCGTTCTTCACCCTTAATCTGTTCATCGGTGTCATCATCGACAACTTCAACGAGCAGA
  AGAAGAAAGCCGGTGGCTCGCTGGAAATGTTCATGACGGAGGATCAGAAAAAGGTTCC

- 1532I/1534C:

  GGAACCAGGTGGGCAAGCAGCCAATTCGCGAGACCAACATCTACATGTACCTCTACTTCGTGTTCTTCA
  **T**CATCT**G**CGGGTCGTTCTTCACCCTTAATCTGTTCATCGGTGTCATCATCGACAACTTCAACGAGCAGA
  AGAAGAAAGCCGGTGGCTCGCTGGAAATGTTCATGACGGAGGATCAGAAAAAGGTTCC

- 1532T/1534C:

  GGAACCAGGTGGGCAAGCAGCCAATTCGCGAGACCAACATCTACATGTACCTCTACTTCGTGTTCTTCA
  **C**CATCT**G**CGGGTCGTTCTTCACCCTTAATCTGTTCATCGGTGTCATCATCGACAACTTCAACGAGCAGA
  AGAAGAAAGCCGGTGGCTCGCTGGAAATGTTCATGACGGAGGATCAGAAAAAGGTTCC
