## Supplementary material for "Novel ddPCR diagnostic assays for the high-throughput surveillance of *Kdr* mutations in *Aedes albopictus*": S1 Table

### Supplementary file 6: Cost estimations of traditional genotyping tools compared to the novel ddPCR TaqMan assays

| Approach | Cost type | Personnel time (h) | Unit cost (€) | Total cost (€) |
|---|---|---|---|---|
| | 96-wells DNA extractions (CTAB) + QC check (Qubit) | | 1.5 | |
| Individual genotyping by | PCR (AmpliTaq Gold) + Gel QC check | | 1.6 | 12200 |
| Sanger sequencing | Sanger sequencing (subcontracting) | | 3.0 | |
| (20 populations × 100 specimens) | Personnel (96 samples/day × 20 days) | 140 | 21.5 | 3010 |
| | | | **Total** | **15210** |
| | 96-wells DNA extractions (CTAB) + QC check (Qubit) | | 1.5 | |
| Individual genotyping by | Taqman qPCR (Biorad CFX 96 qPCR system) | | 1.5 | 6000 |
| TaqMan qPCR | Reagents cost per sample | | 3.0 | |
| (20 populations × 100 specimens) | Personnel (96 samples/day × 20 days) | 140 | 21.5 | 3010 |
| | | | **Total** | **9010** |
| | 96-wells DNA extractions (CTAB) + QC check (Qubit) | | 1.5 | |
| Individual genotyping by | PCR + barcoding + QC + pooling (in house) | | 3.0 | 9200 |
| mass multiplexed sequencing | Sequencing as 2x150 bp reads to 15x (subcontracting) | | 0.1 | |
| (20 populations × 100 specimens) | Personnel (96 samples/day × 20 days + 4 days data analysis) | 168 | 21.5 | 3612 |
| | | | **Total** | **12812** |
| | Pooled gDNA extractions (CTAB) + QC check (Qbit) | | 2.2 | 138 |
| TaqMan ddPCR on pools | TaqMan ddPCR (Biorad QX200 ddPCR system) | | 4.7 | |
| (20 pools of 100 specimens) | Personnel (20 samples in 2 days) | 14 | 21.5 | 301 |
| | | | **Total** | **439** |

Cost and time-to-results were estimated for the screening of 1 Kdr locus from 20 *Ae. albopictus* populations each comprised of 100 mosquito specimens. Unit costs are calculated per DNA sample or per working hour for personnel. Personnel time was estimated from mosquito sample reception to final data analysis. The cost of lab equipment required to perform the work is not included.
